# Spike development inhibition in the *ftin* mutant is associated with multiple phenotypes and regulated by multiple biological pathways

**DOI:** 10.1101/2020.10.13.338731

**Authors:** Yongsheng Zheng, Jinpeng Zhang, Cheng Liu, Han Zhang, Xiajie Ji, Mumu Wang, Hui wang, Rongzhi Zhang, Ruyu Li, Weihua Liu

**Affiliations:** Institute of Crop Sciences, National Key Facilities for Crop Gene Resources and Genetic Improvement, Chinese Academy of Agricultural Sciences, Beijing 100081, P. R. China; Crop Research Institute, Shandong Academy of Agricultural Sciences, Jinan 250100, P. R. China

**Author notes:** Co-first author (Yongsheng Zheng, Jinpeng Zhang). Corresponding author: Professor Weihua Liu, Institute of Crop Sciences, Chinese Academy of Agricultural Sciences, Beijing 100081, P. R. China,; Additional corresponding author: Professor Ru-Yu Li, Crop Research Institute, Shandong Academy of Agricultural Sciences, Jinan 250100, P. R. China.

**Keywords:** Bread wheat, Spike-development inhibition, Cold Response, Proteomics, Indoleacetic Acids, RNA Interference

## Abstract

Spike development of wheat line 3558M was strongly inhibited by low temperature stress in spring. The *fertile tiller inhibition* (*ftin*) gene in the wheat line 3558M is associated with multiple phenotypes, including the production of fewer tillers, delayed floral transition, and death of the shoot apical meristem. We systematically investigated the genes and pathways underlying the differences using ITRAQ proteomics and RNA-sequencing technologies and found multiple biological pathways including to the cold acclimation pathway and multiple defence responses (e.g. reactive oxygen species-mediated hypersensitive response, salicylic acid-mediated systemic acquired resistance) are activated and led to tillers death of the wheat line 3558M under cold stress. Meanwhile, the cold acclimation pathway inhibited the SVP-SCO1-LFY flowering pathway and led to delayed floral transition. Particularly, two TaPIN proteins were significantly downregulated, and multiple auxin signalling genes were also differentially expressed. Knocking down the two *TaPIN* genes using RNAi technology significantly reduced the tiller number. The cold stress inhibited the auxin transport to reduce the tillers of 3558M. Taken together, the *ftin* gene might be a cold-sensitive mutation and that is the cause of multiple biological pathways and phenotypic changes.

## Introduction

Based on the period development theory, spike development in wheat is divided into three main physiological stages which were tillering stage, the floral transition from vegetative stage to reproductive stage and inflorescence development stage (Sreenivasulu & Schnurbusch, 2012). Tillering is a two-step process: The formation of axillary buds and the outgrowth of the buds. The dormancy of axillary buds is the result of complex interactions between endogenous developmental signals and environmental factors (Kebrom et al, 2012). Bud outgrowth is inhibited by apical dominance and regulated by multiple hormone signalling pathways, e.g., cytokinin (CK), auxin and strigolactones and their interactions (Thimann & Skoog, 1933; Tanaka et al, 2006). The regulation of these two steps, which is dependent on both genotypes and environment, determines the number of productive tillers that form, and thus also the final spike number.

The transition from vegetative to reproductive stage is a critical stage of development in which the outgrowth of tillers will complete the floral transition and finally develop into spikes although not all wheat tillers can develop into spikes (Deng et al., 2011; Qin et al., 2015). In bread wheat, the initiation of the floral transition is mainly controlled by photoperiod (*Ppd*) response, vernalization (*Vrn*), and earliness *per se* (*Eps*) genes (Herndl et al., 2008). The photoperiod response is mainly regulated by *PPD1*, a member of the *PSEUDO-RESPONSE REGULATOR* (*PRR*) gene family, which upregulate the expression of the *VRN3*/*FLOWERING LOCUS T1* (*FT1*) as an activator of the floral transition under long days. Three dominant genes, *Ppd-A1*, *Ppd-B1* and *Ppd-D1*, which are located on wheat chromosomes 2A, 2B and 2D, respectively, determine photoperiod insensitivity (Law, Sutka, Worland, 1978; Scarth & Law, 1983). The wheat carrying the photoperiod-insensitive genotype is rapid flowering regardless of whether they are exposed to short-day or long-day illumination. The vernalization response is mainly regulated by *Vernalization1* (*VRN1*), *Vernalization2* (*VRN2*) and *Vernalization3* (*VRN3*). *VRN1* is positively involved in promoting the floral transition which encodes an APETALA1/FRUITFULL-like MADS box transcription factor. In allohexaploid bread wheat, most of the variation in the vernalization process is controlled by the three homologs *VRN1* locus (*Vrn-A1*, *Vrn-B1* and *Vrn-D1*) that are located on chromosomes 5A, 5B and 5D, respectively (Danyluk, 2003; Yan et al., 2003). *VRN2* encodes a flowering repressor which suppresses the expression of VRN1. *VRN3* encodes a mobile protein that is homologous to *Arabidopsis FT* and is transported from the leaves to the SAM to form a protein complex that regulates the expression of the floral meristem identity gene *VRN1* (Li & Dubcovsky, 2008), *Eps* genes are important for the fine-tuning of flowering and the adaptation of wheat to different environments when the photoperiod and vernalization requirements have been fulfilled. Several *Eps* loci have been mapped to different chromosomes. *LUX ARRHYTHMO/PHYTOCLOCK 1* (*LUX/PCL1*), a candidate gene for *Eps-3*^*Am*^, has been mapped and cloned (Gawroński & Schnurbusch, 2012; Gawroński et al., 2014).

Low temperature is an important environmental stress factor that affects the spike development of wheat. Especially in the spring, when the vernalization has been completed and the spike begin the transition from vegetative stage to reproductive stage, the ability of response to cold stress will dramatically descend and the spike will suffer frost injury to lead to significant yield loss under cold stress. However, how the low temperature affects the spike development and how the wheat responds to low temperature stress is still unclear. For this report, we characterized a wheat mutant line 3558M, which displays multiple phenotypic differences, including a reduced tiller number, a delayed floral transition, large spikes, significantly increased grain number and grain weight, compared with the near isogenic line (NIL) wheat 3558. These phenotypes of the 3558M were controlled by a single recessive gene *fertile tiller inhibition gene* (*ftin*), which was mapped to 1AS and linked closely to markers Xcfa2153 with 1.0 cM genetic distance (Zhang et al., 2013). In the present study, we find that the mutant line 3558M is cold sensitive and cannot response to normal cold stress in spring. To reveal which biological pathways are responsible for the multiple phenotypes of 3558M and might be regulated by the *ftin* gene, we systematically studied the mutant line 3558M using proteomic, transcriptome sequencing and RNAi technologies. Multiple biological pathways were identified that are closely related to the spike development inhibition phenotype. Our results provide the basis for further studies on the regulatory functions of the *ftin* gene and spike development in bread wheat.

## Results

### Multiple phenotypes were significant different between wheat line 3558M and NIL 3558 under low temperature environment

The tillers and spike number of 3558M are significant lower than NIL 3558. Statistical analysis showed that the number of tillers produced before winter without low temperature stress was not significantly different between 3558M and NIL3558 (Fig. 1A). However, at the regeneration stage during a cold environment, 3558M produced tillers significantly fewer than NIL 3558; 3558M produced only approximately two tillers, but 3558 produced approximately seven tillers. Almost all of the tillers of 3558 eventually developed into spikes (average of 10.1), but the tillers of 3558M, which were only produced before winter, eventually developed into spikes (average of 3.2).

**Fig.1.**
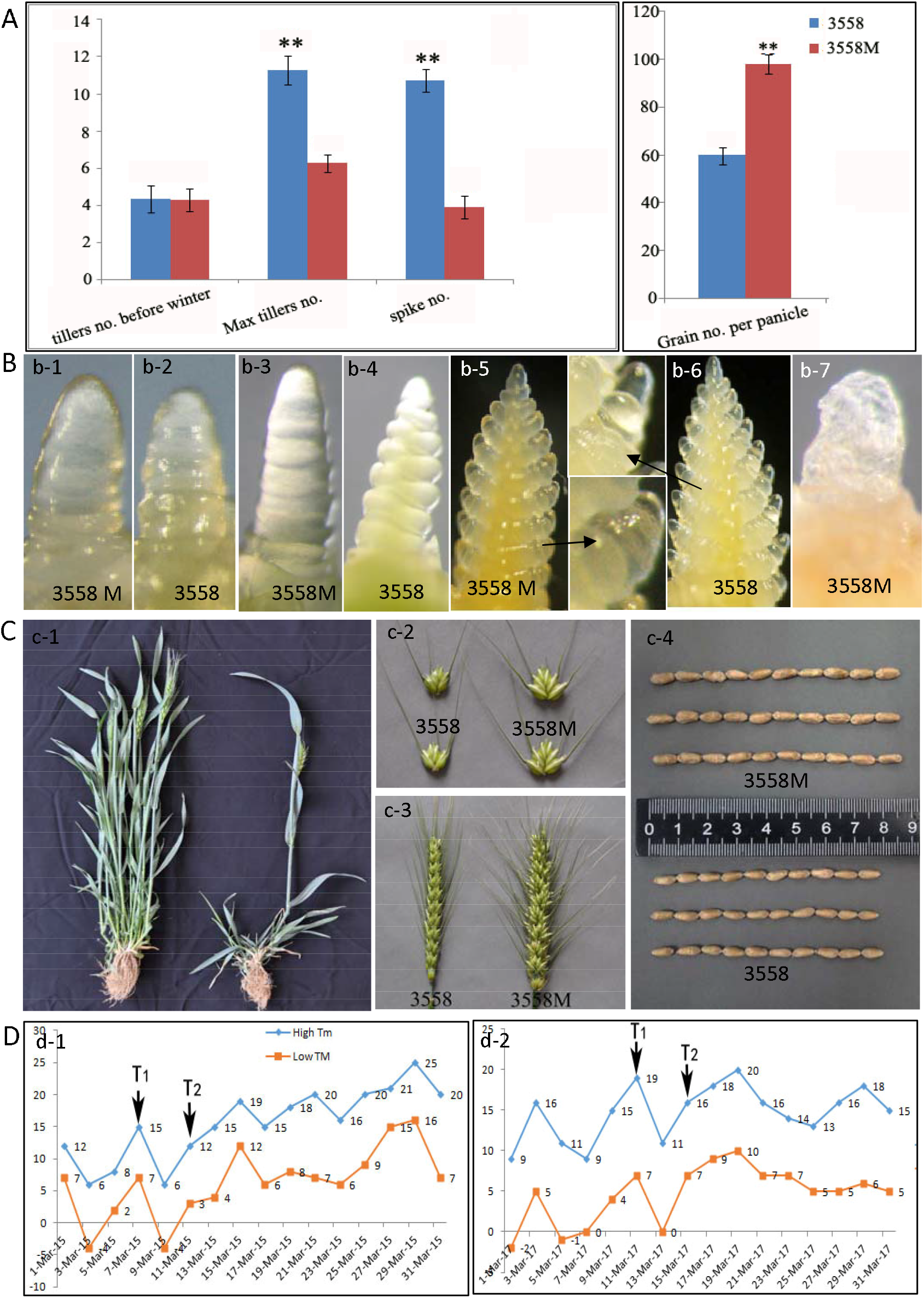
Multiple phenotypic differences between wheat 3558M and NIL line 3558. A. Statistical analysis of the differences in number of spikes, tillers, and grains per spike between 3558MU and 3558 (**stands for p<0.01). B. Anatomical analysis of the differences of SAM in several important physiological stages of single ride (b1 and b2), double ridge (b3 and b4), spikelets differentiation (b5 and b6). b-7, the death of 3558M meristem. C. At heading time, 3558MU had significantly fewer spikes and more grains per spike compared with 3558. D The time points at which samples were taken for RNA sequencing and protein identification (between the two time points T1 and T2, there was a significant cooling change about zero degrees centigrade.

The development process of wheat 3558M is remarkably slower than NIL 3558 at floral transition stage. Anatomical examination of 3558M tillers indicated that the shoot apical meristem (SAM) of tillers at single-ridge stage before winter is not significantly different between the 3558M and 3558 (Fig. 1B). However, the spike rachis of 3558 tillers had begun to develop to the spikelets at double-ridge stage after winter, and the SAM of 3558M tillers has still stay at the single ridge stage (Fig. 1B). When the florets of the spikelets of 3558 had begun to develop pistils and stamens, the spikelets of 3558M are still stay in the florets differentiation stage without the pistils and stamens (Fig. 1B). It’s obvious that the development process of 3558M is significantly slowly than those of the 3558 after winter. In particular, the SAMs of tillers produced after winters not only develop slower, but did not complete the floral transition to develop into spikes. Meanwhile, we found at the floral transition stage, if there are temperature drop fluctuations with normal cold stress (under the zero-degree, Fig 1D), the SAM of 3558M tillers will suffer the freezing damage and lead to death of the shoot apical meristem (SAM) of about two-thirds tillers produced before winter (Fig. 1B and C). Finally, the few tillers of 3558M can develop into spikes.

Although the spike number of 3558M was significantly less, the grain number per spike was significantly more than NIL 3558 (Fig. 1A and C). Generally, one spikelet of the spike of 3558M can produce six to seven grains, and the spikelet of the spike of 3558 can produce three to four grains. Finally, the grain number per spike of 3558M is significant higher than 3558. Meanwhile, the length and weight of grains of 3558M is also significantly higher than NIL 3558. The spike development inhibition phenotype was clearly associated with multiple phenotypic characteristics, including reduced tiller and spike numbers, delayed spike development, tiller apex death, high grain number and weight.

### Determination of hormone and reactive oxygen species contents

To reveal the differences in hormone contents between the SAM of 3558M and 3558 in response to cold stress, we used an LC-ESI-MS/MS system to analyse six types of hormones: auxin, CKs, gibberellins (GAs), abscisic acid (ABA), jasmonates (JAs) and salicylic acid (SA) (Fig. 2). There were no significant differences in the contents of GAs (GA1, GA3, GA4 and GA7) or ABA between the tillers of 3558M and 3558 (Fig. 2A). The JA and N-[(-)-JASMONOYL]-(L)-ISOLEUCINE (JA-L-ILE) contents of 3558M were significantly higher than those of 3558, especially after cold stress (P<0.05, Fig. 2A and B). The IAA content of 3558M was slightly lower than that of 3558, and was decreased in both 3558M and 3558 after cold stress (Fig. 2B). The content of the CK *trans*-Zeatin was significantly higher in 3558M than in 3558, and dramatically increased in both 3558 and 3558M after cold stress (Fig. 2B). The SA content was the highest of all of the detected hormones, reaching over 2000 ng/g in 3558M (Fig. 2C). After cold stress, the SA content was higher in 3558M, but slightly lower in 3558. In particular, although there was no significant difference in the whole SAM between 3558M and 3558, the IAA content of the base of SAM in 3558M is significant lower than in 3558 after cold stress (Fig.2E, *P*<*0.05*).

**Fig. 2.**
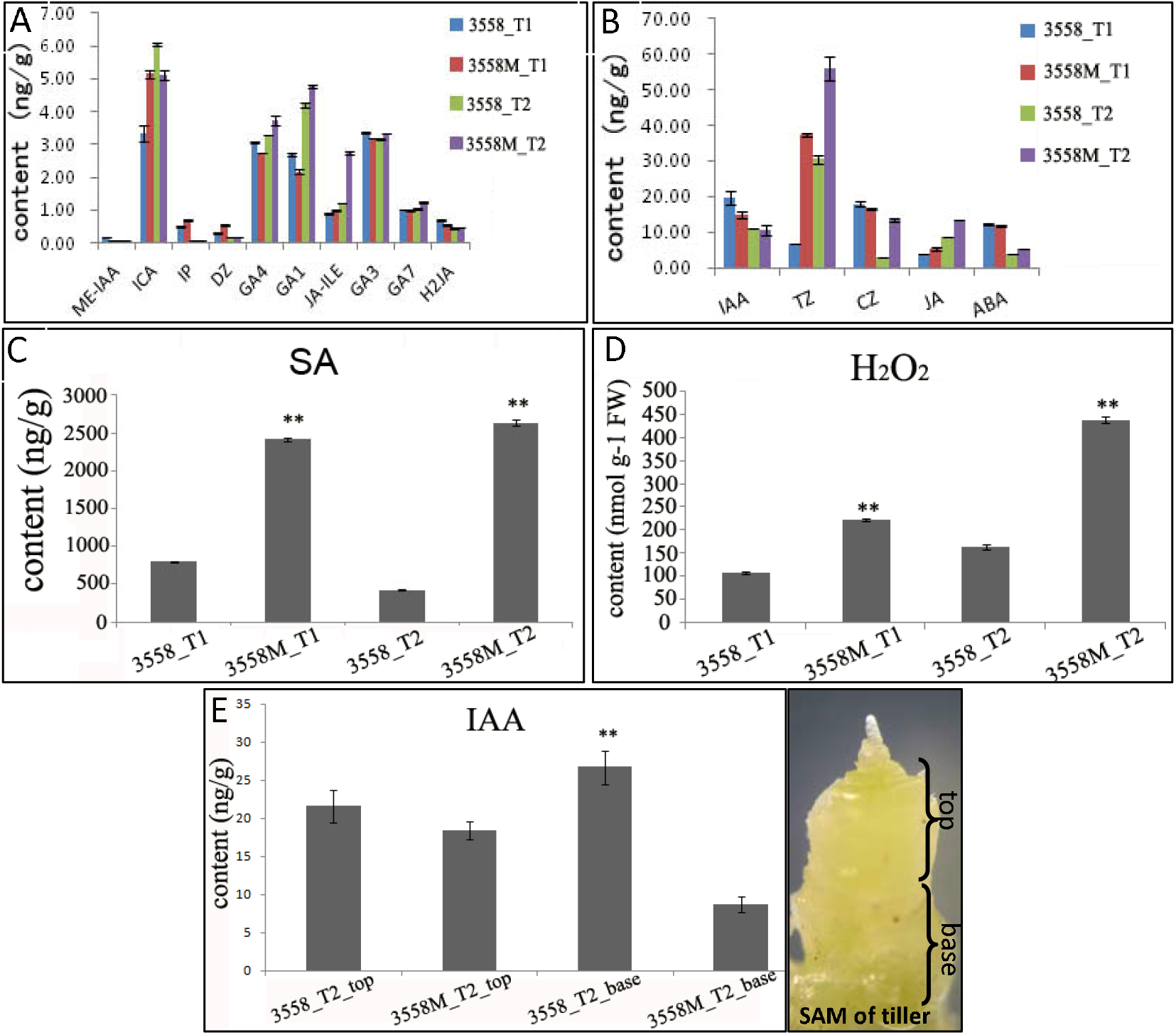
Measurement of hormone and H_2_O_2_ (a reactive oxygen species) contents of 3558 and 3558M (**stands for p<0.01). A and B. The contents of hormones present at concentrations below 10 ng/g (A) and between 10 and 100 ng/g (D). C and D. The contents of salicylic acid (C) and H_2_O_2._ (D). E. The contents of IAA in top and base SAMs of 3558 and 3558M after cold stress.

To further identify the reactive oxygen species (ROS) mediated hypersensitive response and death of the SAM of tillers of 3558M, we further measured the H2O2 concentrations of 3558M and 3558 (Fig. 2D). The pattern of ROS accumulation was similar to that of SA accumulation. ROS levels were significantly higher in 3558M than in 3558. It is worth noting that the ROS concentration of 3558M reached its maximum value after cold stress, but the ROS concentration of 3558 did not change much after cold stress.

### Identification and functional classification of differentially expressed proteins

To reveal the effects of normal cold stress on protein expression in 3558M, we performed a high-throughput quantitative proteomic analysis of 3558M and 3558 using ITRAQ technology. In total, 13,577 proteins (unique peptides ≥2 and spectral counts ≥3) and 9179 proteins were identified in both experimental replicates (Repeat 1 and Repeat 2) (Table S1). Proteins were identified as differentially expressed proteins (DEPs) using strict cut-off criteria (fold change >1.35 or <0.7). Before cold stress, the abundance of 364 proteins in 3558M wheat was significantly different compared with that in 3558; among them, 237 proteins were significantly upregulated, and 175 proteins were significantly downregulated. After cold stress, 611 proteins significantly changed in abundance in 3558M; among them, 335 proteins were upregulated, and 175 proteins were downregulated (Table S2).

To identify more DEPs and upstream regulators, we isolated the nucleus and extracted nuclear proteins from 3558M and 3558 after cold stress to analyse the nuclear proteomics. A total of 9182 nuclear proteins were quantified (unique peptides ≥2 and spectral counts ≥3) (Table S1). Among them, 2976 proteins were only identified in the nuclear proteome. A total of 290 proteins significantly changed in abundance in response to cold stress. Of the 201 upregulated proteins, all were identified as being upregulated under cold stress in the total proteome. The abundances of 58 proteins were significantly different between the total and nuclear proteomes. Of the 89 downregulated proteins, 25 proteins were also identified as being downregulated under cold stress in the total proteome (Table S2). Finally, we performed functional classification and gene ontology (GO) analyses of the DEPs from the nuclear proteome and the total proteome (Fig. 3, P<0.05). Before cold stress, almost all of the upregulated DEPs were involved in photosynthesis, and the downregulated biological pathways mainly included the response to hypoxia and toxin catabolic process. After cold stress, the upregulated biological pathways mainly included the innate immune response (systemic acquired resistance, defence response/incompatible interaction and plant-type hypersensitive response), chitin catabolic process, toxin catabolic process, response to toxic substances and photosynthesis; the downregulated biological pathways mainly included the regulation of chromatin silencing, DNA methylation and DNA packaging. It is worth noting that most of the DEPs in the innate immune response pathway were upregulated in 3558M after cold stress, but they did not significantly different between 3558M and 3558 before cold stress.

**Fig. 3.**
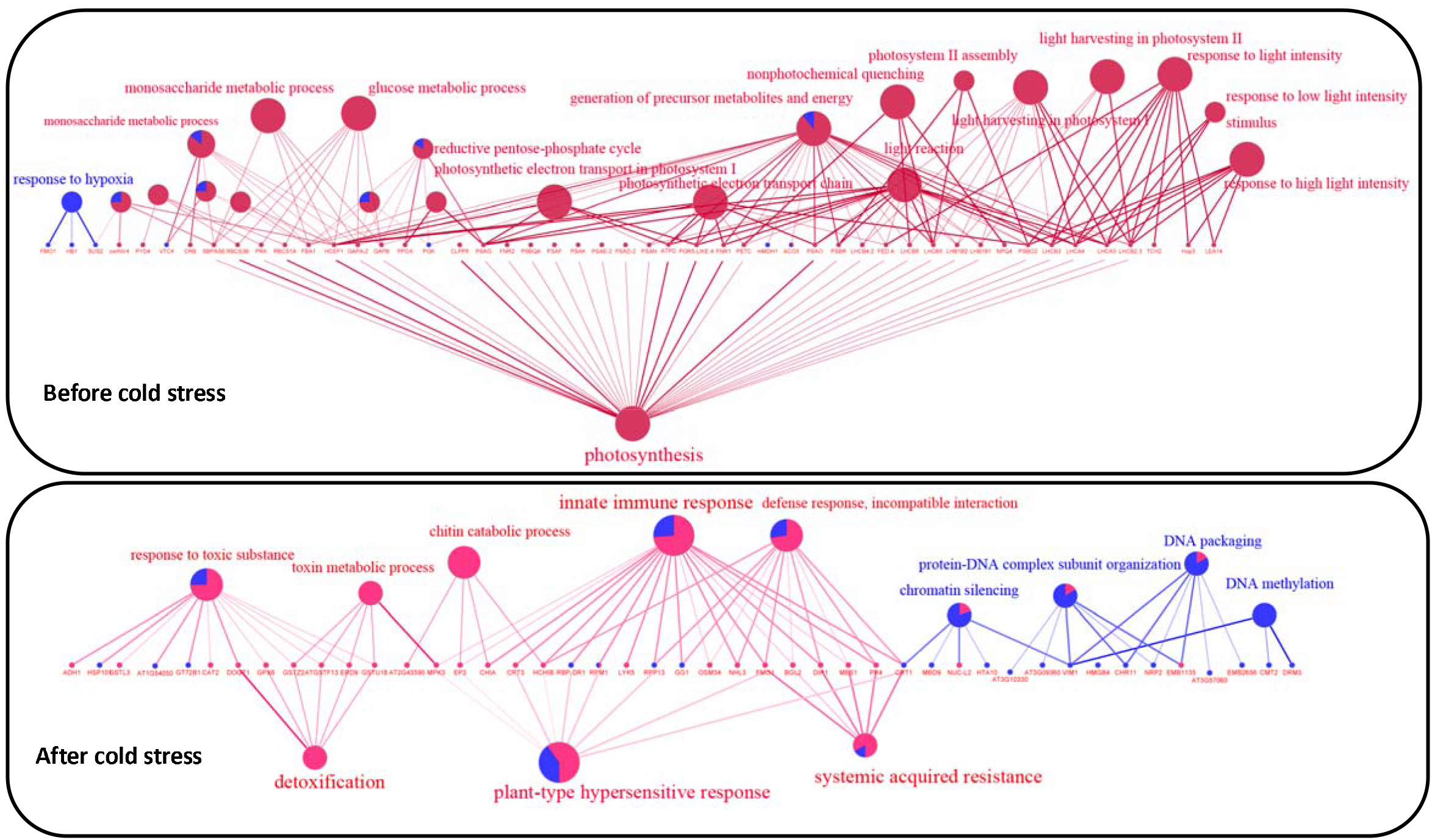
Enriched GO Terms (P<0.05) for differentially expressed proteins of wheat 3558M and 3558WT before and after cold stress. The red stand for upregulated DEPs and the blue stand for downregulated DEPs

### Transcriptome sequencing and identification of differentially expressed genes

Although we identified the top 15,000 proteins in our analysis of the total proteome and the nuclear proteome, only a few upstream regulators were identified. For example, only a few DEPs were predicted as transcription factors (TFs), and only 36 DEPs were identified as TFs. To reveal the differences in upstream regulator dynamics between 3558M and 3558, we performed RNA-seq on the SAMs of 3558M and 3558 before and after cold stress using the Illumina HiSeq 2500 platform. Twelve RNA-seq libraries (four samples with three biological replicates per sample) were prepared, and total of 997,215,482 clean paired-end 150-bp reads, consisting of 149.57 Gb, were obtained with an average GC content of 52.6% (Table S3.1). Among the clean reads, 893,955,370 (89.6%) could be uniquely mapped to the wheat reference genome (Table S3.2). More than 90% of the reads mapped to annotated genes and 88% mapped to protein-coding genes. We detected 48,421 to 50,092 expressed genes with FPKM > 1 in each sample (Table S3.3). Finally, we analysed the differentially expressed genes (DEGs) between 3558M and 3558 according to the criteria |log2Fold Change (FC)| >1 and Q-value < 0.05. In total, we identified 3780 DEGs between 3558M and 3558 before cold stress and 3426 DEGs after cold stress. Before cold stress, 2692 DEGs were upregulated and 1088 DEGs were downregulated in 3558M. After cold stress, 2965 DEGs were upregulated, and 511 DEGs were downregulated in 3558M (Fig 4A and 4B, Table S4).

**Fig. 4.**
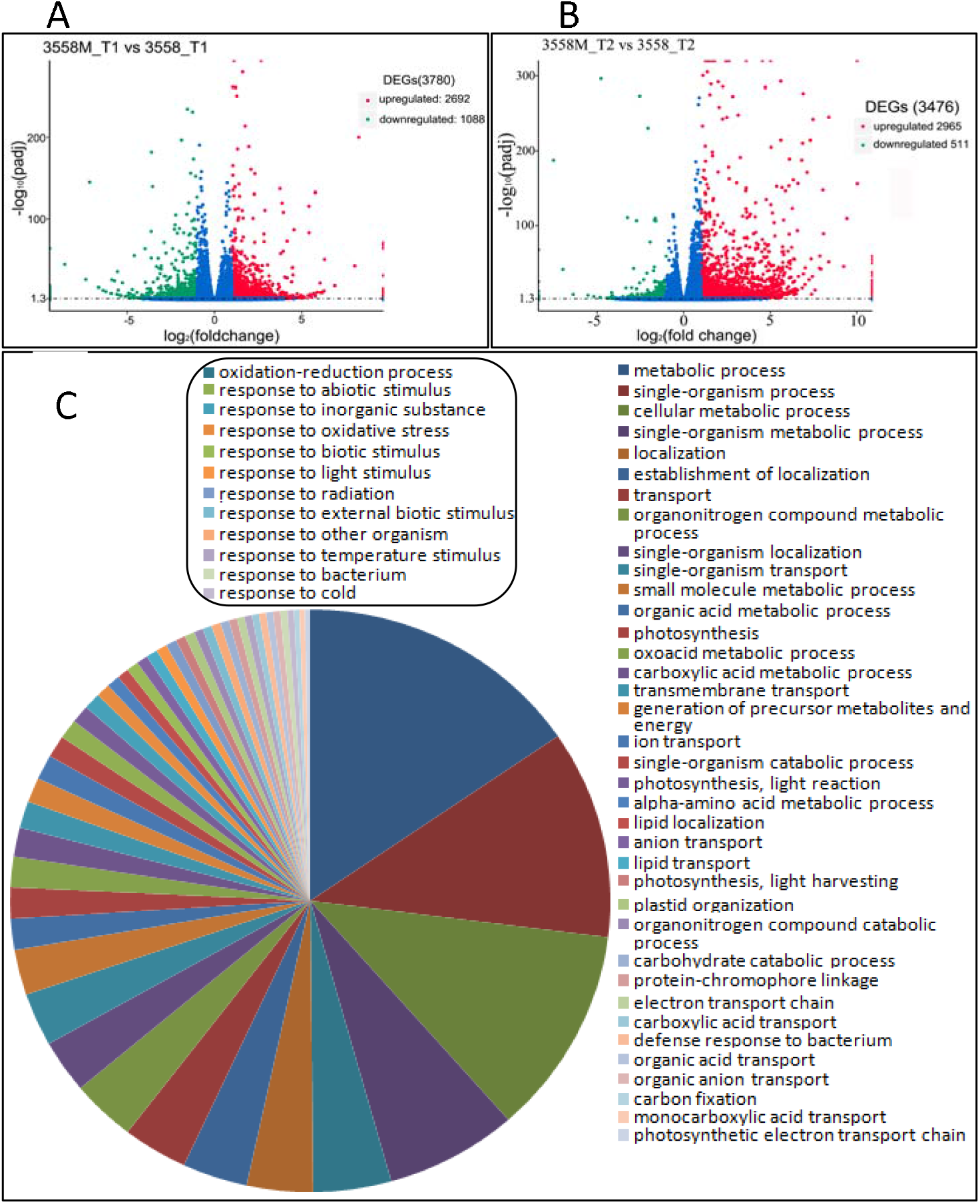
Volcano plot of DEGs and enriched GO Terms (P<0.05) for wheat 3558MU and 3558WT. A and B. The volcano plots showing the number of DEGs before cold stress (A) and after cold stress (B). The red points stand for upregulated DEGs, and green points stand for downregulated. . C. Enriched GO biological process terms for wheat 3558M_T2 and 3558_T2 after cold stress.

To identify the major functional profiles of DEGs, GO enrichment analysis was carried out separately for four sets of DEGs (3558M_T1 vs 3558_T1_up, 3558M_T1 vs 3558_T1_down, 3558M_T2 vs 3558_T2_up and 3558M_T1 vs 3558_T1_down). For the 3558M_T2 vs 3558_T2_up set, 49 GO enriched terms in the biological process category (P<0.05 and DEGs ≥10) were identified; these terms included metabolic process, oxidation-reduction, photosynthesis, and lipid transport (Fig 4C). In particular, 12 significantly enriched GO terms were associated with the response to cold stress, such as oxidative stress, response to cold stress, and biotic and abiotic stimulus, were identified after cold stress (Fig. 4C). The DEGs associated with these terms had the highest expression levels in 3558M_T2. For example, of the 63 DEGs annotated to the GO term oxidation-reduction, 40 showed highest expression levels in 3558MU_T2 (Fig. 5A). Interestingly, few significantly enriched GO terms were identified for the other three sets, and no GO terms associated with response to oxidative stress or cold stress were identified (Table S5).

**Fig. 5.**
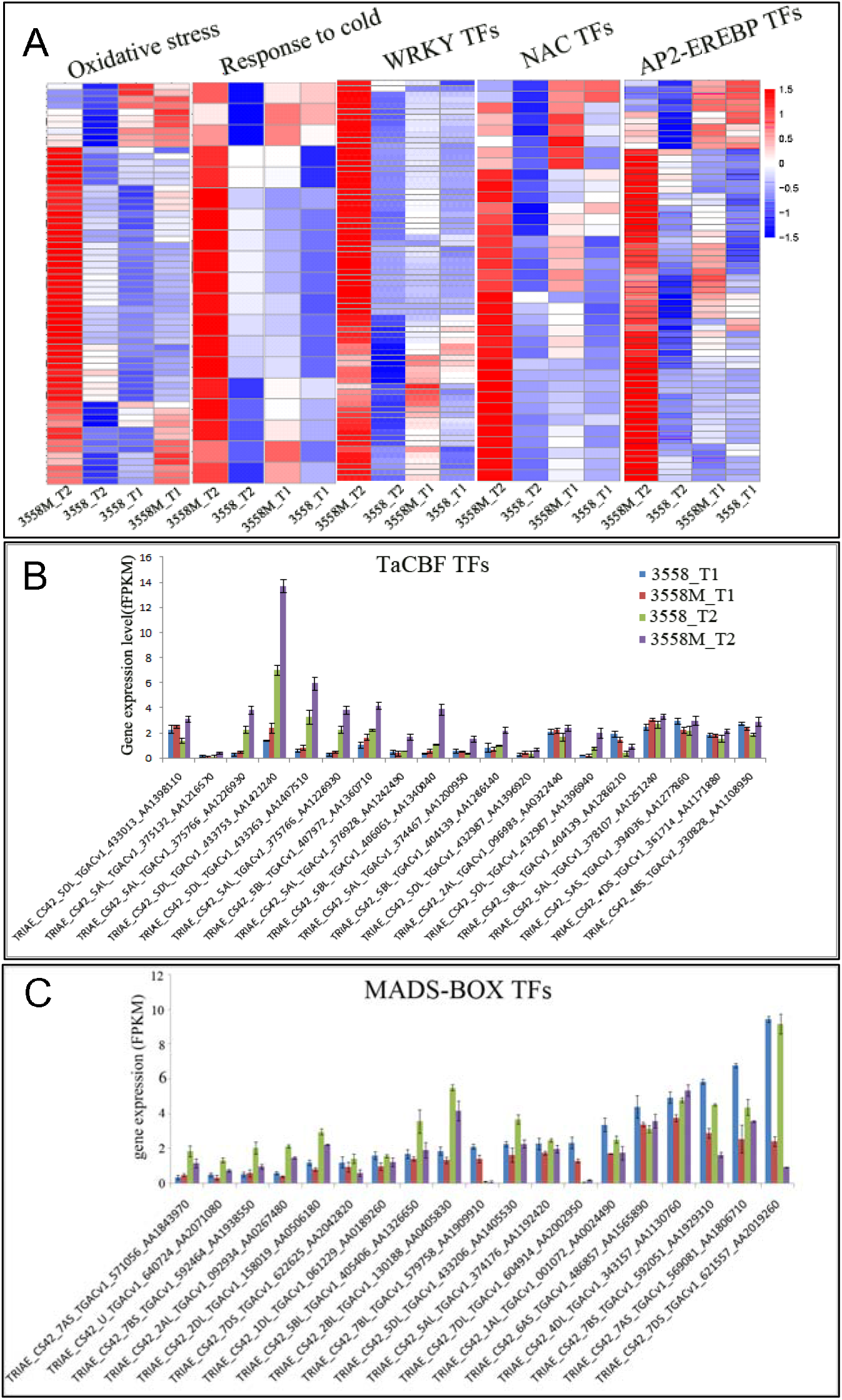
Heatmaps and the gene expression levels of DEPs with different functions. A. Heatmap of the DEPs involved in the biological processes of response to oxidative stress, cold stress and DEPs annotated as TFs containing different functional domains (WRKY, NAC and AP2-EREBP). The red stand for upregulated DEGs, and blue stand for downregulated DEGs. B. The gene expression levels of TaCBF TFs, which are involved in cold stress response. C. The gene expression levels of TF DEPs that contain the MADS-box domain, which ared involved in flowering development.

TFs are important upstream regulators that regulate developmental and defence processes in response to physiological signals and different types of stress. A total of 3292 genes were predicted to encode TFs using iTAK software in PlnTFDB. Among them, 1055 TFs were differentially expressed in at least one pairwise combination (Table S6). A total of 511 TFs were identified before cold stress, and 689 TFs were identified after cold stress. Before cold stress, 227 TFs were significantly upregulated and 284 TFs were significantly downregulated in 3558M compared with 3558. After cold stress, 302 TFs were significantly upregulated in 3558M, and 387 TFs were downregulated. Interestingly, TF DEGs with the same functional domain tended to have the same expression patterns, especially after cold stress. Most of the TF DEGs involved in stress and defence responses showed significantly upregulated expression in response to cold. The upregulated TFs mostly belonged to five main domain families: APETALA2/ethylene-responsive element binding protein (AP2-EREBP) (52), bHLH (23), MYB (23), WRKY (32), and NAC (15). All of the WRKY family members were significantly upregulated in 3558M, and 34 WRKY family members were significantly upregulated in 3558M after cold stress (Fig. 5A). Most AP2-EREBP and NAC family members showed the highest expression level in 3558M (Fig. 5A). Most of the 15 C-repeat binding factor (CBF) TFs, which are members of the AP2-EREBP family, were upregulated in response to cold stress (Fig. 5B). These results imply that multiple defence mechanisms were quickly activated in response to cold stress in 3558M. In contrast, some TF DEGs involved in the positive regulation of flowering time and development were downregulated. The downregulated TFs belonged to six main categories of domain families: TCP (20), MADS-box (11), GRF (11), SBP-box (12), GATA (8), and AUXIN RESPONSE FACTOR (ARF) (4). Both before and after cold stress, 17 of the 19 MADS-box family members were downregulated in 3558M (Fig. 5C).

### The tillers number is significantly reduced after knocking down the two TaPIN genes in wheat using RNAi technology

In the proteome and transcriptome data, multiple DEPs and DEGs were involved in the AUXIN signal transduction. Among them, there were two TaPIN proteins significantly downregulated in 3558M (Table S2). Meanwhile, there was a significant difference in transcription levels between 3558M and 3558 after cold stress (*P*<*0.05*, Fig 6A). To further verify the expression patterns of the two PIN proteins, we used a western blot assay to determine the abundances of these proteins in 3558M and 3558. The results showed that both PIN proteins were significantly downregulated in the wheat line 3558M (Fig. 6B and C), and were consistent with our ITRAQ results. To determine the relevance of the correlation between the spike number and TaPIN protein expression, we constructed an RNAi vector targeting *PINa*, pBIOS2043-PINa-RNAi, and one targeting *PINb*, pBIOS2043-PINb-RNAi, separately. The sense and antisense DNA sequences of *TaPINa* (nucleotides 784 to 1036 bp) and *TaPINb* (1013 to 1312 bp) were synthesized using an ABI 3730xl DNA Analyser and inserted into the plasmid pBIOS2043. Six and eight putative transgenic plants were obtained from bombarding 35 and 48 immature cv. Svevo embryos with pBIOS2043-TaPINa and pBIOS2043-TaPINb (RNAi), respectively. A PCR-based assay confirmed the presence of the transgene in three pBIOS2043-TaPINa-RNAi plants and two pBIOS2043-TaPINb-RNAi plants. Homozygous plants in the T_2_ generation we identified using the same PCR-based assay. Finally, two independent homozygous transgenic lines called 1954-1 (RNAi-TaPINa) and 1955-4 (RNAi-TaPINb) that had a reduced tiller phenotype were used for subsequent phenotype analysis and mRNA/protein expression experiments (Fig. 6D and E).

**Fig. 6.**
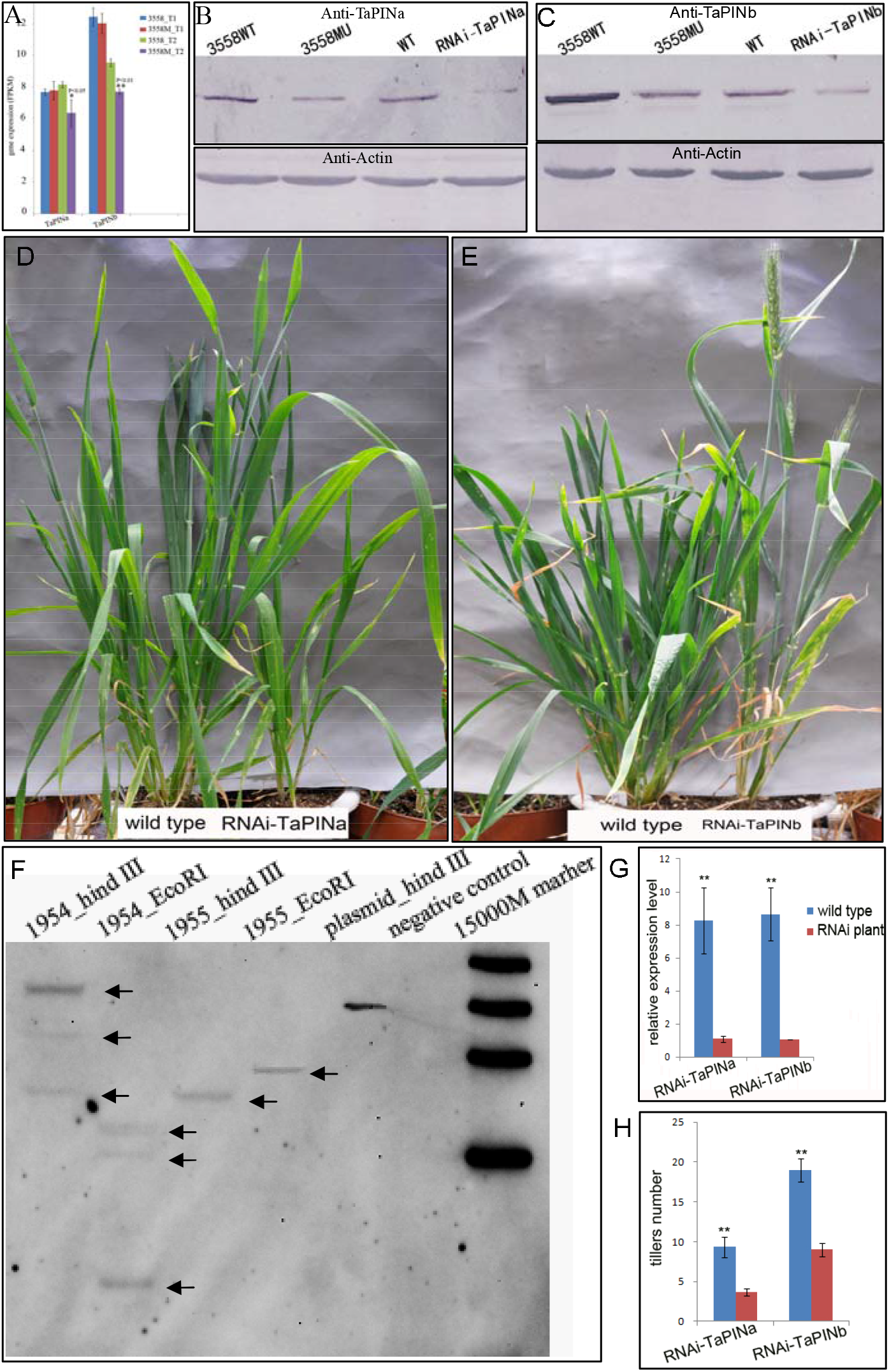
Analysis of TaPINa and TaPINb mRNA and protein accumulation, phenotypic characteristics, and insert number in transgenic RNAi-TaPINa and RNAi-TaPINb plants. A. the genes expression levels of TaPINa and TaPINb in wheat 3558M and 3558; B and C. the proteins expression levels of TaPINa and TaPINb in wheat 3558M and 3558; RNAi-TaPINa, RNAi-TaPINb lines and the wild-type (WT) control, as determined by western blot analysis; D and E. The tiller number phenotype in RNAi-TaPINa, RNAi-TaPINb and WT plants. F. Southern blot analysis revealed three gene copies in RNAi-TaPINa (1954) and one gene copy in RNAi-TaPINb (1955). WT was included as a negative control. G. qRT-PCR analysis of the expression levels of *TaPINa* and *TaPINb* in the RNAi-TaPINa and RNAi-TaPINb lines, respectively, and the WT control (**stands for p<0.01), H. Statistical analysis of the number of tillers and spikes (**stands for p<0.01)

Southern blot analysis suggested that the TaPINa-RNAi and TaPINb-RNAi sequences were integrated into the wheat genome. The 1954-1 line contains three copies of TaPINa-RNAi, and the 1955-4 line contains one copy of TaPINb-RNAi (Fig. 6F). The mRNA and protein abundances of TaPINa and TaPINb were estimated by qRT-PCR and western blot assay, respectively. The qRT-PCR results showed that the abundance of *TaPINa* and *TaPINb* transcripts was reduced by >80% in the TaPINa-RNAi and TaPINb-RNAi lines, respectively (Fig. 6G). The western blot assay results showed that the TaPINa and TaPINb proteins were significantly downregulated in the TaPINa-RNAi and TaPINb-RNAi lines, respectively (Fig. 6A and B). The effect of TaPINa and TaPINb downregulation on the spike number was assessed by comparing the transgenic lines with the wild-type plants. Statistical analysis showed that the axillary buds/tillers of TaPINa-RNAi and TaPINb-RNAi were significantly reduced by >50% (Fig. 6H), but all of the produced tillers eventually developed into spikes. As the spring wheat, the Fielder and transgenic-Fielder did not experience a low temperature stress during their life cycle. So the death of meristem in the TaPIN RNAi lines and delayed flowering time was not observed. On the contrary, the flowering time is earlier in the TaPINb-RNAi line, than in wild type. Finally, the number of tillers is congruent with the number of spikes without aborted phenotype. The results showed that the downregulation of TaPIN proteins results in a reduced spike number phenotype.

## DISCUSSION

### The wheat line 3558M is cold sensitive at the regeneration stage and activates multiple defence responses

Previous studies of inhibition of spike development in wheat line 5660M have shown that multiple cold-regulated (COR) proteins and CBF genes are significantly upregulated and that high expression levels are closely related to the 5660M inhibition phenotype (Zheng et al., 2013). The wheat line 3558M contains a recessive mutation in the same gene mutated in 5660M, *ftin*, and has a similar spike development inhibition phenotype as 5660M. To further verify the cold-sensitive phenotype of the mutant 3558M, we compared the shoot apexes of tillers of 3558M and 3558 before and after cold stress (Fig. 2E and 2F). Eight COR proteins were identified, and all were upregulated in 3558M proteome as determined using ITRAQ in combination with nano LC-MS/MS analysis (Table S2). By analysing the transcriptome of 3558M, most of the CBF/DREB TFs were significantly upregulated after cold stress (Fig 5B). The changes in CBF/DREB TFs expression in 3558M were consistent with those in 5660M according to qRT-PCR analysis (Zheng et al., 2013). Although the *ftin* gene has not been cloned, the fact that the mutant 3558M has the same tiller inhibition phenotype and the same expression trends of CBFs and COR proteins as the mutant 5660M indicates that the recessive mutation of *ftin* in these lines causes cold sensitivity.

GO analysis of DEPs showed that 18 DEPs potentially involved in plant innate immunity were upregulated in 3558M after cold stress. Among these, ten DEPs were involved in the ROS-mediated plant-type HR (Fig. 4). Meanwhile, GO analysis showed that 63 DEGs associated with oxidative stress were significantly upregulated after cold stress (Fig. 5A). ROS are signalling agents that drive the hypersensitive response and are correlated with disease resistance (Asai & Yoshioka, 2010). We found that the relative H_2_O_2_ concentration in 3558M was consistently higher than that in 3558 and reached a maximum value after cold stress (Fig. 2D). Consistent with the higher concentrations of ROS, two NADPH oxidase (RBOH) proteins involved in the generation of ROS were also found to be significantly upregulated after cold stress. RBOH proteins can generate ROS, which antagonize SA-dependent HR signals and suppress the spread of cell death (Torres et al., 2005). FLAVIN-DEPENDENT MONOOXYGENASE1 (FMO1) protein which is involved in defence activation and the HR in *Arabidopsis* (Bartsch et al., 2007) was upregulated after cold stress. VASCULAR ASSOCIATED DEATH1 (VAD1), a regulator of cell death and defence responses, was significantly upregulated after cold stress (Lorrain et al., 2004). Most of the proteins in the hypersensitive response pathway were not significantly different before cold stress but were significantly upregulated after cold stress. In addition, other DEPs involved in the hypersensitive response, such as EP3, CHIN, Calreticulin 3 (CRT3), HCHIB and RPM1, were upregulated after cold stress. At the transcription level, four RBOH and twenty CDPK genes were also significantly upregulated and had the same expression trends as those DEPs after cold stress. Taken together, these results indicate that 3558M experienced a strong ROS-mediated HR under cold stress. The activation of this response may explain why tiller damage and cell death in the SAM occur in 3558M after cold stress.

The results of the hormone content analysis showed that the content of the defence response-related hormone SA in 3558M was significantly higher than that in 3558 (Fig. 2C). Five DEPs involved in SA-mediated SAR were identified, namely FMO1, BGL2, DIR1, SABP2, and PR4. SABP2 is required to convert methyl salicylate to SA as part of signal transduction pathways that activate SAR in systemic tissue (Forouhar et al., 2005). The SABP2 protein was identified in both the nuclear proteome and the total proteome and was significantly upregulated after cold stress. The ankyrin-repeat protein NPR1 and WRKY TFs are the central positive regulators of SAR, transducing the SA signal to activate PR gene expression (Li et al., 1999). Five of six NPR1 genes were upregulated after cold stress at the transcriptional level. Three homeologs of NPR1-type protein 2 were significantly upregulated after cold stress. Multiple WRKY TFs have been shown to be involved in the SA-mediated SAR pathway. WRKY71 functions as a regulatory factor upstream of NPR1 and PR1b (Liu et al., 2007). Although NPR1 and WRKY TF proteins were not detected by ITRAQ, seven pathogenesis-related (PR) proteins downstream of the SAR pathway were identified and found to be significantly upregulated: four PR1 proteins, one PRB1-3, one PRB1.2, and one PR10. In summary, the ROS-mediated HR and SA-mediated SAR were strongly activated after cold stress in 3558M, indicating that the normal cold stress induced multiple forms of plant innate immunity in 3558M.

### Flowering genes and their gene regulatory networks were downregulated overall, resulting in delayed spike development in 3558M

Phenotypic analysis of 3558M revealed that spike development was inhibited and delayed in spring. Analysis of proteome and transcription data revealed many spike development genes, most of which were downregulated in 3558M, consistent with the inhibited spike development phenotype of 3558M. MADS-box TFs, such as *OsMADS14*, *OsMADS22*, *OsMADS34*, *OsMADS4*, *OsMADS50*, *OsMADS55,* and *OsMADS58*, are involved in the regulation of flowering time and the transition from the inflorescence meristem to the spikelet meristem phase. In the present study, 19 MADS-box TFs showed downregulated expression after cold stress (P<0.05, Fig. 5C). The *SOC1* gene can repress the expression of *FRIGIDA* (*FRI*) and integrate signals from the photoperiod, vernalization and autonomous floral induction pathways in *Arabidopsis* (Liu et al., 2008; Lee et al., 2000). The three *TaSOC1* genes, which are homeologous MADS-box TFs located in the wheat A, B and D subgenomes, encode proteins sharing 86% and 62% amino acid identity with SOC1 in rice and *Arabidopsis*, respectively. All three genes were significantly downregulated in 3558M compared with 3558 both before and after cold stress. The LEAFY (LFY) protein promotes early establishment of floral meristem identity by directly activating the *APETALA1*, *CAULIFLOWER* and *AGAMOUS* genes. Interestingly, all three *LFY* homeologs, which encode proteins sharing 83% and 64.8% identity with LFY in rice and *Arabidopsis*, respectively, were significantly downregulated in 3558M after cold stress. Three homeologs of *VRN-1* were also downregulated in 3558M. In particular, TaAGL10 and TaAGL11AP1 TFs, were downregulated at the protein level. Sixteen SBP-box TFs were differentially expressed between 3558 and 3558M, 11 of which were downregulated in 3558M at the transcription level after cold stress. One SBP transcription factor, Squamosa promoter-binding-like protein 13 (SPL13), was found to be downregulated at the protein level after cold stress. SPL proteins can specifically bind to the consensus nucleotide sequence 5’-TNCGTACAA-3’ in MADS-box TF promoters to activate expression and promote both vegetative phase change and flowering. Previous studies of 5660M have shown that, as in 3558M, genes involved in spike development and the floral transition (e.g., *VRN1* and *LFY*) are downregulated in 5660M (Zheng et al., 2013). The results indicate that the block or delay in spike development in 3558M and 5660M after exposure cold temperatures and is closely related to the later flowering of 3558M and 5660M tillers.

### The reduced tillers phenotype is closely associated with the downregulation of PIN proteins

In the present study, we found that the contents of auxin in both 3558 and 3558M were significantly reduced after cold stress. Although the total contents of auxin in the whole SAMs organization of 3558M and 3558 were not significant differently, the auxin content in the top SAM organization of 3558M was significantly higher than that in the base SAM organization in 3558 (Fig. 2E). We speculate that auxin transport is inhibited and maintained at a low level at the base organization of the tillers where the lateral buds/tillers grow out. This may be an important cause of the production of fewer tillers. Multiple DEPs and DEGs were annotated as being involved in auxin signal transduction and significant downregulated in 3558M after cold stress. Two PIN proteins were found to be downregulated in the proteome of 3558M and western blot assays of the TaPIN proteins TaPIN1a and TaPIN1b showed that both proteins were significantly downregulated in 3558M compared with 3558, which was consistent with the results of the ITRAQ analysis. One aminopeptidase M1 protein (APM1), which can bind to the auxin transport inhibitor N-1-naphthylphthalamic acid and plays a negative role in the regulation of PIN auxin transport proteins (Murphy et al., 2006), was significantly upregulated in 3558M compared with 3558. Significant upregulation under cold stress was observed for one TOPLESS (TPL) protein, which functions as a co-repressor with AUXIN/INDOLE-3-ACETIC ACID (AUX/IAA) transcriptional repressors to bind ARF proteins and repress the expression of auxin response genes (Krogan et al., 2012; Szemenyei et al., 2008). There were 64 DEGs related to auxin signalling: 10 AUX1 genes, 22 AUX/IAA genes, 7 ARF genes, 19 SAUR genes and 6 GH3 genes (P<0.05). Particularly, all of the seven ARF genes were significantly downregulated after cold stress. Taken together, these results show that the auxin signal transduction pathway might be influenced by cold stress and also suggest that the reduced tillers phenotype of 3558M could be related to auxin transport. When we used RNAi technology to knock down the *TaPINa* and *TaPINb* genes, spike number was significantly reduced. However, the heading and maturation dates of the TaPINb-RNAi (1954) lines were significantly earlier than those in the wild type, and there were no significant differences in heading between TaPINa-RNAi (1955) lines and wild type. It is thus clear that downregulation of TaPINa and TaPINb did not delay spike development. Therefore, the delayed spike development and reduced spike number phenotypes were associated with a different biological pathway(s).

### The *ftin* recessive gene in 3558M influences multiple biological pathways to produce multiple phenotypes changes

Tiller formation has been intensely studied in rice and wheat. For example, *OsMOC1*, which controls tillering and tiller formation, has been mapped to the long arm of chromosome 6 and encodes a plant-specific GRAS transcription factor (Li et al., 2003). In bread wheat, the recessive gene *tin* inhibits tiller formation and has been mapped to the short arm of chromosome 1AS (Hyles et al., 2017). The candidate gene for *tin* has been cloned and encodes a cellulose synthase-like protein with homology with members of the CsIA clade (Hyles et al., 2017). We have performed related research on tiller formation and spike development in our lab. In a previous study, we mapped the *ftin* gene from a wheat–*Agropyron cristatum* (L.) Gaertn.-derived line 3558M to 1AS and found that it was closely linked (1.0 cM distance) to marker Xcfa2153 (Zhang et al., 2013). The *tin* gene is also closely linked to marker Xcfa2153 (Hyles et al., 2017). To further determine whether the two genes are the same locus, we analysed variation of the *tin* gene and found that the *tin* gene sequence was the same between 3558M and 3558 (unpublished data). This indicates that *ftin* and *tin* are likely not the same gene, although they cause a similar low tiller number phenotype. In this report, we found that the *ftin* gene led to multiple phenotypic differences between 3558M and 3558 wheat. The *ftin* gene not only altered the tiller number but also delayed spike development and caused sensitivity to normal cold stress. There were multiple other phenotypes (e.g., large spikes with high grain no. and weight) at maturity. Systematically investigation of the differences between 3558M and 3558 using proteomic, RNA-seq and RNAi technology provided insight into which metabolic pathways are potentially regulated by the *ftin* gene.

First, the wheat line 3558M is cold sensitive and not response to normal cold stress at the regeneration stage. The ICE-CBF-COR signal cascade system was intensely active to response to normal cold stress in both 3558M and 5660M wheat. After extreme cold stress, the ROS-mediated HR and defence response in 3558M wheat were activated and led to the death of the apexes of 3558M tillers. As a result of the apex death, sections of tillers did not develop into spikes. In response to normal cold stress, multiple innate immune responses (e.g., the HR-, SAR- and FLS2-induced pathogen-associated defence systems) were activated in 3558M wheat.

Second, consistent with the known role of *OsPIN* genes in the regulation of tiller development in (Chen et al., 2011), two downregulated TaPIN proteins and multiple DEGs involved in the auxin signalling pathway were identified. Moreover, all of the ARFs, which positively regulate the auxin signalling pathway, were downregulated after cold stress, and all of the *TaGH3* genes, which negatively regulate the auxin signalling pathway, were significantly upregulated (Staswick et al., 2005). The auxin content was also reduced after cold stress, especially in the SAM of 3558M. These findings suggest that mutant *ftin* gene influence the auxin signalling pathway and that cause the reduced tiller phenotype of 3558M. This notion is further supported by the finding that knockdown of *PIN* gene expression causes a reduced tiller phenotype. A previous study showed that the regulator OsNPR1, which regulates the SAR pathway, can affect rice growth and development by disrupting the auxin pathway at least partially through indirectly upregulating OsGH3.8 expression (Li et al., 2016). In the present study, we identified six *TaNPR1* genes at the transcription level, five of which were significantly upregulated after cold stress. Furthermore, the death of SAM caused by ROS-mediated HR affects the synthesis and distribution of auxin. Taken together, our results suggest that the reduced tillering phenotype of 3558M is caused by a disrupted auxin signalling pathway that might result from the activation of the SAR pathway and multiple active defence responses.

Third, the delayed floral transition is associated with the downregulation of flowering network genes. Using our data, we attempted to elucidate the mechanism by which the *ftin* gene interferes with/regulates the flowering pathway to delay the floral transition of 3558M. The floral transition in wheat is regulated by the three types of genes vernalization (*Vrn*), photoperiod (*Ppd*) response, and earliness *per se* (*EPS*) and their interactions with temperature. In the present study, we found that the expression of the clock regulators *PRR5* and *PRR7*, and flowering genes that play key roles in photoperiodic flowering responses, *GI*, *CRY*, and *FT*, was slightly upregulated in 3558M (Table S7.1). We also compared the expression of the *Vrn* genes, including repressors and activators. In our unpublished data, we identified 18 MADS-box TFs as repressors that were silenced by vernalization and 9 MADS-box TFs as activators induced when vernalization is complete and the floral transition is initiated. Among the 18 repressors, 11 MADS-box TFs were completely silenced in 3558M; there was no significant change in the expression of the other 7 MADS-box TFs (Table S7.2). We also compared the expression of the *VRN2* (*ZCCT1 and ZCCT2*) genes and found they were completely silenced in both 3558M and 3558. These results show that delayed floral transition is likely not caused by alteration of the photoperiodic or vernalization pathway. We identified three *TaSOC1* gene homeologs and three *LFY* homeologs, which are downstream targets of SOC1. Both sets of homeologs, which positively regulate spike development, were significantly upregulated. In *Arabidopsis*, crosstalk between the cold response and flowering pathways is mediated by *SOC1* (Seo et al, 2009). It has been reported that SOC1 can directly bind to the promoters of *CBF* genes and repress their expression. In *Arabidopsis*, SVP, another flowering repressor, plays an important role in the response of plants to ambient temperature changes (Lee et al., 2007) and associates with the promoter region of *SOC1* to directly repress its transcription in the shoot apex. In the present study, we found that both alleles of *TaSVP* were significantly upregulated in 3558M, especially after cold stress. The expression patterns of *TaSVP*, *TaSOC1* and *TaLFY* genes in 3558M under cold stress are consistent with its delayed spike-development phenotype. Therefore, we speculate that after cold stress, *TaSVP* senses the ambient temperature change and represses the expression of *TaSCO1* and *TaLFY*. The *ftin* gene in 3558M delays spike development through the SVP-SOC1-LFY thermosensory pathway.

To sum up, although we have not cloned the *ftin* mutant gene, according to the current data, the *ftin* mutant gene might be a cold-sensitive mutation and that is the cause of multiple metabolic pathways and phenotypic changes.

## Materials and methods

### Plant materials

The wheat mutant line 3558M (T. *aestivum* L., 2n = 6x = 42, AABBDD) and the NIL 3558 (T. *aestivum* L., 2n = 6x = 42, AABBDD) are near isogenic lines obtained from the seventh generation derived from the cross Fukuhokomugi (T. *aestivum* L., 2n = 6x = 42, AABBDD) /A. *cristatum* Z559 (A. *cristatum* (L.) *Gaertn*., 2n = 4x = 28, PPPP; National Gene Bank accession number Z559)//Youla (T. *aestivum* L., 2n = 6x = 42, AABBDD) (Zhang et al., 2013). Both of them are winter wheat and planted on the three plots in Jinan on October7, 2014 and October10, 2016 in autumn at above 20 °C. Each plot contained 24 rows, 2 m in length, and separated by 0.25 m. Crop management was according to the local practice. Both of them would go through a long winter and the tillers will be stop to product as the temperature usually falls below zero. In the spring of next year, the tillers will be going on tillering as the temperature rises. Based on the period development theory, the spike differentiation of winter wheat is divided into several important physiological stages: single ride stage (vegetative growth), double ridge stage (floral transition), spikelets differentiation stage, florets differentiation (pistils and stamens). The SAM of 3558M and 3558 were harvested at the double ridge stage before and after a normal cold stress when the temperature has a zero degree fluctuations (Fig 1.D). Finally, approximately 3 g SAM of 3558M and 3558 were collected and stored at −80°C in a cryogenic refrigerator to extract protein, hormone and RNA. The common spring wheat cultivar Fielder is used to transgenic experiments. The seeds of Fielder were grown in a greenhouse at 25 °C with a photoperiod of 16 h light and 8 h dark for 7–10 d. The leafs of Fielder (Wild type) and transgenic Fielder (1954-1 and 1955-4) at the double ridge stage were harvested to quantitative RT-PCR and Southern blotting analysis. As the spring wheat, the Fielder and transgenic Fielder did not experience a low temperature stress during their life cycle.

### Methods

#### Determination of plant hormone content

The SAMs of twelve samples (Table S8, two varieties × three biological replicates × two time points, ~100 mg each) were harvested from the tillers of 3558MU and 3558 at T1 (on Mar. 11, 2017) and T2 (on Mar. 15, 2017), immediately frozen in liquid nitrogen, and ground into powder. Hormone extraction and ESI-Q TRAP-MS/MS analysis were performed according to the protocol of Wuhan Metware Biotechnology Co., Ltd.

### Proteomic analysis

#### protein extraction and labeled with iTRAQ 4plex

About the total proteome, the shoot apical meristem of eight wheat samples (Table S8, two varieties ×2 biological replicates × two time points ~100mg each) were harvested from the tillers of the 3558MU and 3558 at the T1 (on Mar. 7, 2015) and T2 (on Mar. 14, 2015).

The SAM of tillers of wheat 3558M and 3558 were ground into powders in liquid nitrogen, and the powders were each suspended overnight in 30 mL of a 10% w/v TCA/acetone solution that contained 0.07% v/v mercaptoethanol at 20◻. After centrifugation at 40 000 × g for 1 h at 4C, the pellets were each washed with 30 mL of ice-cold acetone. After vacuum drying, pellets were reconstituted in 900ul 8M guanidine and pH was adjusted to >7.5 with 1M Tris HCl pH8.1. Disulfide bond was reduced by addition of 20 mM DTT and incubated at 60◻ for 30 min. After cooling to room temperature, 40 mM iodoacetamide was added and the samples were kept in the dark for 1 hour to block free cysteine. Protein assay was done at this step with Bradfrod method. The samples were dialyzed against 2 M urea and 50 mM NH4HCO3 for 2 hours and 50 mM NH4HCO3 for 2 hours twice. 200ug of protein was digested by Trypsin at 1:50 (w:w, trypsin:sample) and incubated at 37°C overnight. The digested peptides were further filtered with Amicon ultra centrifugal filters (10kD MWKO, Merck Millipore, MA), dried under vacuum extensively to remove residual primary amine from reagents. ½ of the digested samples were labeled with iTRAQ 4plex reagent (ABSciex, MA, US) according manufacturer’s instructions and combined after labeling and dried. About the nuclear proteome, four wheat samples (two varieties ×2 biological replicates ~500mg each) were harvested from the tillers of the 3558MU and 3558 at the T2 (on Mar. 14, 2015). The proteins were extracted and then labeled with iTRAQ 4plex reagent according to above methods.

#### MS identification

First dimensional separation of TMT labeled peptides High-pH Reverse phase HPLC were used for peptide fractionation using Gilson 300 series. The iTRAQ4plex labeled and combined samples were solubilized in 200 μl of 20 mM ammonium formate (pH10), and injected onto an Xbridge column (Waters, C18 3.5 μm 2.1X150 mm) using a linear gradient of 1%B/min from 2-45% of B (buffer A: 20 mM ammonium formate, pH 10, B: 20 mM ammonium formate in 90% acetonitrile, pH10). 1 min fractions were collected and speed vac dried. 2.2.2 Nano LC-MSMS Selected fractions were analyzed by nanoLC-MS/MS using a RSLC system interfaced with a Q Exactive™ Hybrid Quadrupole-Orbitrap Mass Spectrometer (ThermoFisher, San Jose, CA). Samples were loaded onto a self-packed 100 μm x 2cm trap packed with Magic C18AQ, 5μm 200 A (Michrom Bioresources Inc, Aubum, CA) and washed with Buffer A(0.2% formic acid) for 5 min with flow rate of 10ul/min. The trap was brought in-line with the homemade analytical column (Magic C18AQ, 3μm 200 A, 75 μm x 50cm) and peptides fractionated at 300 nL/min with a multi-stepped gradient (4 to 15% Buffer B (0.16% formic acid 80% acetonitrile) in 35 min and 15-25%B in 65 min and 25-50%B in 55 min). Mass spectrometry data was acquired using a data-dependent acquisition procedure with a cyclic series of a full scan acquired in Orbitrap with a resolution of 120,000 followed by MSMS scans (30% of collision energy in the HCD cell) of 20 most intense ions with a repeat count of one and the dynamic exclusion duration of 30 sec.

#### Databases Searching

The LC-MSMS peak list of 13 fractions from each experiment were searched in MUDPIT style against the corresponding Uniprot database (Uniprot_Triticum aestivum (Wheat)_20150401, 94854 sequence) using an in house version of X!tandem (SLEDGEHAMMER (2013.09.01), the gpm.org) with carbamidomethylation on cysteine and iTRAQ label on lysine and N-terminus of peptides as fixed modification and oxidation of methionine as variable modification. A +/−5 ppm and 20 ppm were used as tolerance for precursor and product ions respectively. Intensity of iTRAQ4plex reporter ions of each spectrum was extracted using an in-house perl script and corrected for isotope cross-over using values supplied by the manufacturer. The False Discovery Rate was estimated for all samples by using a reverse database (FDR<0.01). The treatment/blank control ratio of each spectrum was calculated using reporter ion intensity and normalized to the median ratio of all identified spectra that fit certain criteria: peptide belongs to homo sapiens database, peptide log (E)<=−2 (E value<=0.01), total reporter ion intensity >40,000 for each pair. Spectra that have ratio as “Divide 0” were replaced with an arbitrary number “10”. Pairwise ratios (115/114 and 117/116) of individual protein were calculated using median ratio of all peptides belonging to the protein that has at least two data points that fit the criteria of peptide log€ <−2 and sum of reporter ions of both channel >20,000.

### Transcriptome analysis

RNA isolation, cDNA library construction and sequencing: the shoot apical meristem of 12 wheat samples (Table S8, two varieties ×three biological replicates × two time points ~100mg each) were harvested from the tillers of the 3558MU and 3558 at the T1 (on Mar. 11, 2017) and T2 (on Mar. 15, 2017). Total RNA was extracted using RNeasy Plant Mini Kit (TIANGEN, DP432) according to the manufacturer’s protocol. RNA purity was checked using the NanoPhotometer® spectrophotometer (IMPLEN, CA, USA). RNA concentration was measured using Qubit® RNA Assay Kit in Qubit® 2.0 Flurometer (Life Technologies, CA, USA). RNA integrity was assessed using the RNA Nano 6000 Assay Kit of the Bioanalyzer 2100 system (Agilent Technologies, CA, USA). The sequencing libraries preparation and high-throughput sequencing were carried out by Novogene Bioinformatics Technology CO., Ltd. (Beijing, China, http://www.novogene.com).

Western blot assay and quantitative RT-PCR: The protein extracts were from the SAMs of wheat 3558MU_T2, 3558_T2, Fielder wheat and their transgenic lines 1954-1 and 1955-4 (Table S8). The western blot assays and quantitative RT-PCR were performed as described by Zheng et al. (2013 & 2017). Specific primers for TaPINa and TaPINb (TaPINa-F: GTGGACCATCACGCTCTTCT, TaPINa-R: GAGCATGAGCGTGTACCAGA; TaPINb-F: CTCTCTCGGACAAGGAAACG, TaPINb-R: AGCAGGCTTGGTGAAGAAGA) were used to detect expression levels. The relative expression levels were calculated using the relative 2−ΔΔCt method.

RNA interference and Southern blotting analysis: The conserved coding sequence of *TaPINa* (784 to 1036 bp) and *TaPINb* (1013 to 1312 bp) were selected separately as the RNAi target sequence and used to design a RNAi hairpin structure that was inserted into pBIOS2043. After sequencing the target sites, the binary vector was transformed into the wheat cultivar Fielder by *Agrobacterium tumefaciens*-mediated transformation. The gene-specific primers 5’-ATTCTTATTTCTTTCCAGTAGC and 5’-AGAAGCGGCATAATGTGAGA were used to amplify a 478-bp fragment of the FAD2 intron, a fragment of the interference vector. Genomic DNA was extracted from the leaves of T3 transgenic plants (Table S8, lines 1954-1 and 1955-3) using a plant genomic DNA Extraction Kit (Tiangen Biotech, Beijing, China). About 20 μg of DNA was successfully digested with 5 U of EcoRV and incubated at 37 °C for 24 h. The digested genomic DNA fragments were separated on a 0.8% (w/v) agarose gel, and transferred onto Zeta-Probe GT nylon membrane (Bio-Rad, Hercules, CA, USA). The DNA fragments were fixed to the membrane by UV cross linking. The 1000-bp PCR fragment of the FAD2 intron was labeled with digoxin. Probe labeling, hybridization, washing and detection were performed according to instructions of the DIG-High Prime DNA Labeling and Detection Starter Kit II (Roche, Mannheim, Germany).

## Acknowledgements

This research was supported by grants from the National Key Research and Development Program of China (2016YFD0100102), China Agriculture Research System (CARS-03) and Taishan Scholars Project (tsqn201812123).

